# Heterochromatin flexibility contributes to chromosome segregation in the cell nucleus

**DOI:** 10.1101/2020.12.01.403832

**Authors:** Martin Girard, Monica Olvera de la Cruz, John F. Marko, Aykut Erbaş

## Abstract

While there is a prevalent genome organization in eukaryotic cells, with heterochromatin concentrated at the nuclear periphery, anomalous cases do occur. Deviations of chromatin distribution are frequent, for example, upon aging, under malignant diseases, or even naturally in rod cells of nocturnal mammals. Using molecular dynamic simulations, we study the segregation of heterochromatin in the cell nucleus by modeling interphase chromosomes as diblock ring copolymers confined in a rigid spherical shell. In our model, heterochromatin and euchromatin are distinguished by their bending stiffnesses, while an interaction potential between the spherical shell and chromatin is used as a proxy for lamin-associated proteins. Our simulations indicate that in the absence of attractive interactions between the nuclear shell and the chromatin, the majority of heterochromatin segregates towards the nuclear interior due to depletion of less flexible heterochromatin segments from the nuclear periphery. This inverted chromatin distribution is in accord with experimental observations in rod cells. This “inversion” is also found to be independent of the heterochromatin concentration and chromosome number, and is further enhanced by additional attractive interactions between heterochromatin segments. as well as by allowing bond-crossing to emulate topoisomerase activity. The usual chromatin distribution, with heterochromatin at the periphery, can be recovered by further increasing the bending stiffness of heterochromatin segments or by turning on attractive interactions between the nuclear shell and heterochromatin. Overall, our results indicate that bending stiffness of chromatin could be a contributor to chromosome organization along with differential effects of HP1*α*-driven phase segregation and of loop extruders, and interactions with the nuclear envelope and topological constraints.

## INTRODUCTION

In few-micron-sized eukaryotic cell nuclei, meter-long DNA molecules are organized via a dense array of nucleic acid-binding proteins. Histone proteins first forms DNA-protein complexes, which then hierarchically form nucleosomes, the monomers of the heterogeneous chromatin polymer [1]. Depending on the physical proximity and biochemical modifications of nucleosomes, chromatin is either found in a condensed form, heterochromatin, or in a more open form, euchromatin. Heterochromatin is often associated with gene silencing, while euchromatin allows access to lineage-specific genes [2–5]. Consequently, all biological characteristics of a cell are affected by the partitioning of genome into heterochromatin- and euchromatin-rich portions in the nucleus [5–7]. Consistently, many malignant diseases, including carcinoma [8, 9] and progeria [10, 11], as well as aging-related cell dysfunction [12–14], correlate with anomalies in the nuclear organization of chromatin.

In most healthy eukaryotic cells, interphase chromatin with heterochromatin markers segregates towards the surfaces of nuclei (near nucleoli or the inner nuclear envelope (NE) [9, 15–17]. The localization of heterochromatin at the nuclear periphery is usually attributed to attractive interactions between the shell and chromatin provided by proteins such as lamin B receptor (LBR) or lamin A/C proteins [18–20]. In accord with this picture, low expression levels of lamin A/C or LBR results in a depletion of peripheral heterochromatin followed by a coalescence of heterochromatin in the nuclear interior [21–23]. Likewise, such inverted heterochromatin patterns have also been observed in cells that do not express LBR naturally [23, 24]. Consistent with this, loss of peripheral heterochromatin is a hallmark of aging in *Caenorhabditis Elegans* cells upon the redistribution of various nuclear envelope proteins [13]. Findings from various experiments suggest a strong correlation between central heterochromatin coalescence and weak chromatin-NE interactions. Nevertheless, the underlying driving forces, particularly those arising due to the polymer nature of chromatin, remain to be determined.

Understanding behavior of chromatin copolymer *in vivo* is further complicated by the very different chemistry of its euchromatin and heterochromatin sections. For instance, recent experiments have shown that heterochromatin and the HP1 protein undergo a liquid-liquid phase separation, while euchromatin does not [25–28]. Indeed, for polymers, chemical affinity differences between the involving species (heterochromatin and HP1 in this case) can lead to microphase separation instead of homogeneous mixtures, in which domains with various sizes and morphologies can be observed [29, 30]. In addition to that, polymer physics also suggests that inhomogeneous flexibility along a polymer chain can affect the phase behavior as well [31–33]. Indeed, computational and experimental studies show that variations in nucleosome positioning along the chromatin can induce inhomogeneities in chromatin morphology [34–36]. However, how these variations manifest themselves in *in vivo* chromatin flexibility is a rather grey area due to the difficulty of measuring spatial correlations between nearby nucleosome units in their native environment. For the same reason, values measured for the persistence length (the smallest length scale below which polymer behaves like stiff rod, *l*_*p*_, which characterizes the flexibility) of chromatin rather lies in broad range, from *l*_*p*_ ≈ 30 nm to *l*_*p*_ ≈ 300 nm [37, 38]. Nevertheless, in the continuum limit, the persistence length of a fiber with an average thickness of a scales as *l*_*p*_ ~ *a*^4^. This scaling indicates that even a small difference between the effective thicknesses of hetero- and euchromatin portions of the chromatin could provide a considerable difference in the flexibility, and thus, may provide a mechanism for nuclear chromatin organization that might contribute to in concert with direct energetic demixing interactions.

The idea that the difference in mechanical flexibility of heterochromatin and euchromatin can contribute to the nuclear organization via entropic effects is not new. Previously, Monte-Carlo (MC) simulations of relatively short (i.e., *N* = 50 effective monomers) homo-polymer rings of model heterochromatin and euchromatin in spherical confinement showed a concentration-dependent segregation mechanism [39]. Heterochromatin rings were shown to weakly segregate towards the nuclear periphery for volume fractions above 10% to minimize segmental bending energy [39]. However, it is not clear whether and how the nuclear organization changes when the hetero- and euchromatin sections are joined together, as one would expect different behavior of coblock polymers. Furthermore, the interplay between the various parameters (e.g. volume fraction and stiffness differences) and their potential effects on nuclear organization have not been elucidated.

In order to study how mechanical heterogeneity along chromatin polymers affects nuclear organization, we model interphase chromosomes as diblock ring copolymers by using a well-known coarse-grained polymer model [40–43] confined in a spherical shell. The long chromatin chains (i.e., *N* = 1000) are composed of prescribed lengths of heterochromatin and euchromatin blocks, in which the heterochromatin block is always less flexible than euchromatin (Fig. 1A). Our Molecular Dynamics (MD) simulations show that when there is no attraction between the inner surface of the shell and chromatin, the majority of heterochromatin localizes in the nuclear interior by depleting euchromatin, in accord with the experimental observations of heterochromatin inversion [23, 24]. The central heterochromatin localization is observed for any moderate difference in the flexibility (i.e., factor of 2 or less) and does not depend on either chromosome number or heterochromatin content. The inversion increases further if there is a selective attraction between heterochromatin segments. Further, weak attractive interactions (i.e., on the order of thermal energy *K*_*B*_*T*) between heterochromatin and shell can reverse the heterochromatin inversion by depleting heterochromatin from the nuclear interior. The conventional nuclear organization (more heterochromatin at periphery) is also obtained if the persistence length of heterochromatin is on the order of the dimension of the shell.

**FIG. 1.**
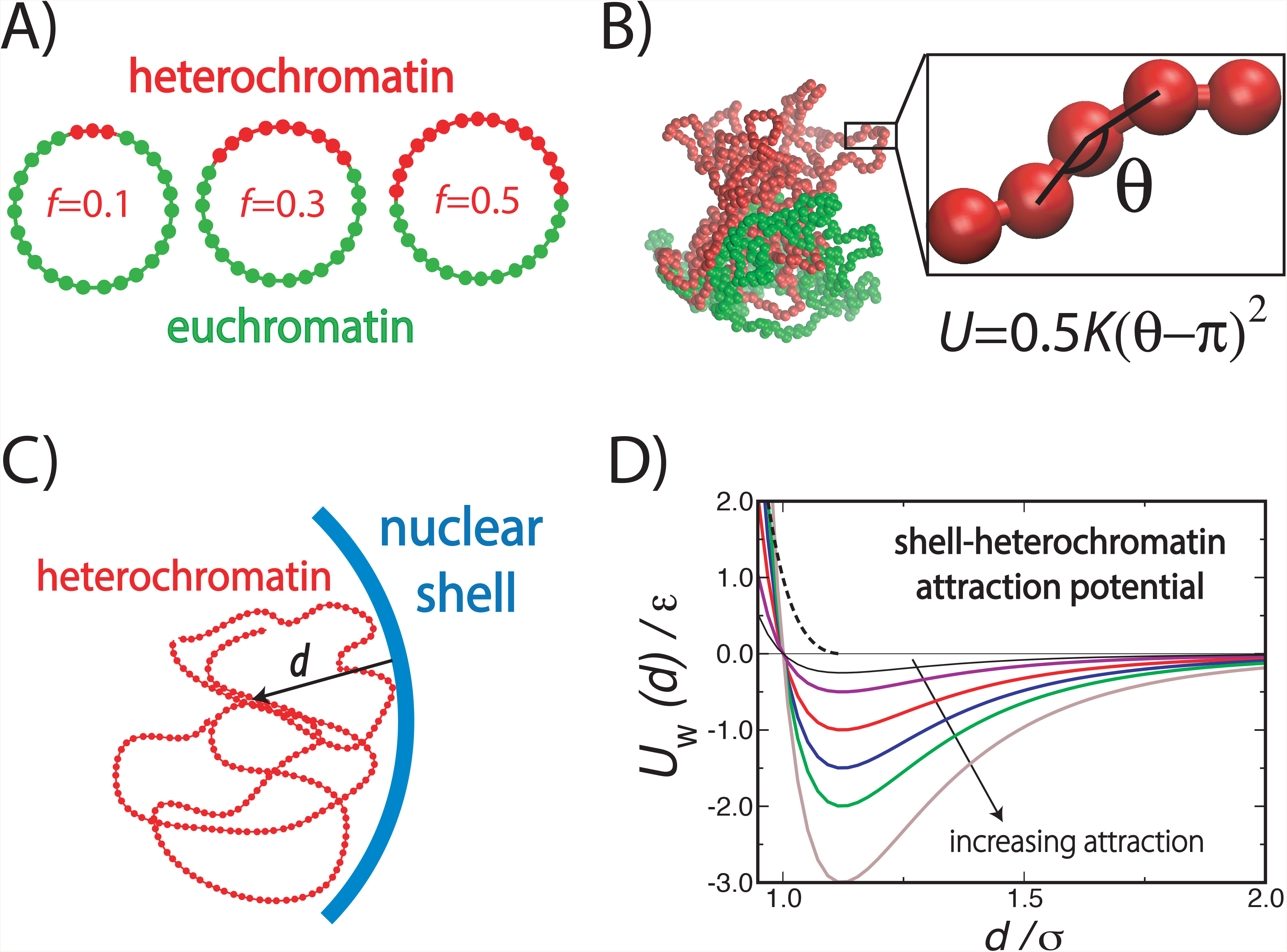
Simulation model. A) The schematics of individual chromatin rings with various fractions of heterochromatin content, *f*. B) An arbitrary simulation snapshot showing an isolated single chromatin chain with *f* = 0.5, and the schematics of the bending potential applied to heterochromatin (red beads) only to alter its persistence length (Eqs. 3, 4). C) The schematics of the attractive potential between heterochromatin beads and the confining shell. D) The plot of the attractive potential, Eq. 1, as a function of then rescaled distance from the shell, *d*/*σ*. The arrow indicates the direction of increasing strength of the potential, *u*/*ε*. The dashed curve is the WCA potential.

## METHODS

In our MD simulations, interphase chromosomes are modeled as bead-spring chains, which is a common model to study large scale and long time behavior of biological and synthetic polymers [43]((Fig. 1A,B). In simulations, *n*_*ch*_ =6-16 unconcatenated copolymers with ring topology are confined in a rigid sphere to mimic nuclear confinement. The ring topology has been shown to model chromosome territories successfully [40, 41, 44]. Each chain is composed of *N* = 1000 beads of size *σ*. Our chromatin model is highly coarse-grained, in which each bead represents roughly 10 nucleosomes resulting in around ≈ 1.6 × 10^6^ basepairs (bps) per chain if we asssume 160 bps per nucelosome and up to 3 × 10^7^ bps in total. The radius of the confining shell is *R* = 20*σ*, which leads to various volume fractions *ϕ* = *n*_*ch*_*Nσ*^3^ /(8*R*^3^) ranging between 10% ≲ *ϕ* ≲ 25%. Thus, our systems may correspond to weak (e.g., yeast) and moderate (e.g., *Drosophila*) confinement levels [42].

All the steric interactions between beads, and beads and confinement are adapted from the 12-6 Lennard-Jonnes (LJ) pairwise potential,

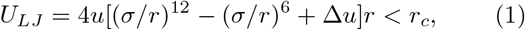

where the strength of the potential is *u* = 1*ε* unless noted otherwise, the energy units in the simulations is *ε* = 1*k*_*B*_*T*, the cut-off length is *r*_*c*_, above which *U*_*LJ*_ = 0, and the shift factor is Δ*u* = 0. To obtain ideal mixture conditions (i.e., the Flory parameter, *χ*_0_ = 0) in an athermal-solvent limit, bead-bead interactions are modeled via the Weeks-Chandler- anderson (WCA) potential by setting *r*_*c*_ = 2^1/6^*σ* and Δ*u* = 1/4 (dashed curve in Fig. 1D) unless noted otherwise.

Interactions between the inner surface of the sphere and all beads were also modeled by above mentioned WCA potential (Fig. 1C) to avoid the diffusion of monomers outside of the sphere. To model the attractive interactions between the heterochromatin and NE, the value of the attraction strength is varied between 0.1*ε* ≤ *u* ≤ 3.0*ε* with *r*_*c*_ = 2.5*σ*(Fig. 1D).

Springs connecting two adjacent chain beads separated by a distance *r* are taken care of by the Finite-Extensible non-extensible (FENE) potentials, which does not allow bond-crossing,

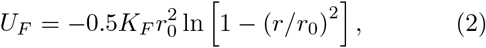

where the bond strength is *K*_*F*_ = 30*ε*/*σ*^2^, and the maximum bond stretch is *r*_0_ = 1.5*σ*. Eq. 2 provides a bond length of *b* ≈ 1*σ*.

To construct our heteropolymers, a harmonic angular potential, which energetically penalizes the bending of heterochromatin segments, is used to control the flexibility of a prescribed fraction, *f*, of each chain (Fig. 1A, B). By varying *f* as 0.1 ≤ *f* ≤ 0.5, various heterochromatin contents are obtained. The form of the angular potential is

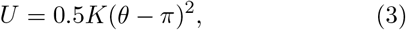

where the spring constant is *K* = 2*ε*/rad^2^ unless noted otherwise, and the angle between three consecutive beads of the corresponding chain segment is *θ* (Fig. 1B). The potential, Eq. 3, is applied only to heterochromatin blocks. For heterochromatin, *K* may be related to *l*_*p*_ as [45],

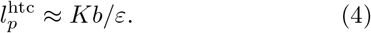

For euchromatin blocks, the persistence length is equivalent to 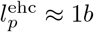.

MD simulations are carried out using the HOOMD-blue molecular dynamics engine [46, 47] with initial configurations built by the Hoobas molecular builder [48]. The simulation boxes are maintained at constant temperature by using a Langevin thermostat with a damping coefficient of *γ* = 1*δt*, where 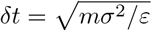 is the LJ time unit with a unit mass of *m* = 1. A simulation time step of Δ*t* = 0.002*δt* is used. Polymer chains are prepared as perfect circles followed by relaxation of a minimum of 10^7^ MD steps. The 10^9^ MD steps (i.e., 2 × 10^6^*δt*) are run for data production. Data is acquired for every 10^4^ MD steps. Visualization is done by using OVITO [49] and VMD [50], while trajectory analyses are done by using custom C++ codes. VMD sphere scale in the snapshots is set to 0.6.

## RESULTS

### Heterochromatin concentration is higher at nuclear center in the absence of chromatin-shell attraction

The experiments on various cell types repeatedly confirmed that in the absence of certain lamin-associated proteins, heterochromatin mostly localizes in the nuclear interior as opposed to the general trend, in which the majority of the heterochromatin resides near the NE [9, 20, 23, 24]. The emergence of this inverted distribution inside the nucleus requires a driving force that can sustain an asymmetric, rather than uniform, heterochromatin distribution. To investigate whether the mechanical heterogeneity along the chromatin polymer can alone provide such segregation by itself, we first consider in our simulations the scenario in which there is no net attraction between the inner surface of the confining spherical shell and the chromatin. In this way, we mimic the experimental conditions where heterochromatin-shell interactions are weak [23]. Since in our simulations heterochromatin and euchromatin monomers interact with the same pairwise potential, Eq. 1, (i.e., all monomers are chemically identical), the equilibrium chromatin organization will mainly be determined by the difference in the bending fluctuations of the two chromatin types.

First, we consider the case in which nuclear concentrations of heterochromatin and euchromatin are equivalent, (i.e. each chromatin chain is composed of an equal number of heterochromatin and euchromatin monomers, *f* = 0.5, see Fig. 1A). Thus, there is no concentration bias towards any of the chromatin types, but heterochromatin has half the flexibility of euchromatin (i.e., *K* = 2 in Eq. 3) [39]. To quantify the chromatin distribution across the nucleus, we calculate the radial concentration profiles for both heterochromatin and euchromatin as a function of the rescaled distance from the nuclear center, *r*/*R*, by using our simulation trajectories (Fig. 2). The concentration profiles rescaled by the total concentration, *ρ*/*ρ*_0_, exhibit a highly non-uniform behavior across the nuclear volume; Near the nuclear center (i.e., *r*/*R* → 0), a significant and systematic increase of the heterochromatin concentration is accompanied by a depletion of euchromatin regardless of the volume fraction (Fig. 2A,B), analogous to the experimentally observed inverted heterochromatin distribution [23, 24]. At the nuclear center, the heterochromatin concentration is almost fifty percent higher than euchromatin (Fig. 2A,B), and this difference diminishes as *R*/*r* → 1 within the error bars. Note that the errors in Fig.2A increase as *R*/*r* → 0 due to the smaller number of monomers near the center. Nevertheless, as compared to an all-euchromatin scenario (i.e., *K* = 0, dashed curves in Fig.2B), our quantitative observations of central heterochromatin coalescence is robust within the simulation time window, which is approximately three times the theoretical relaxation time of a linear chain with equal length [51].

**FIG. 2.**
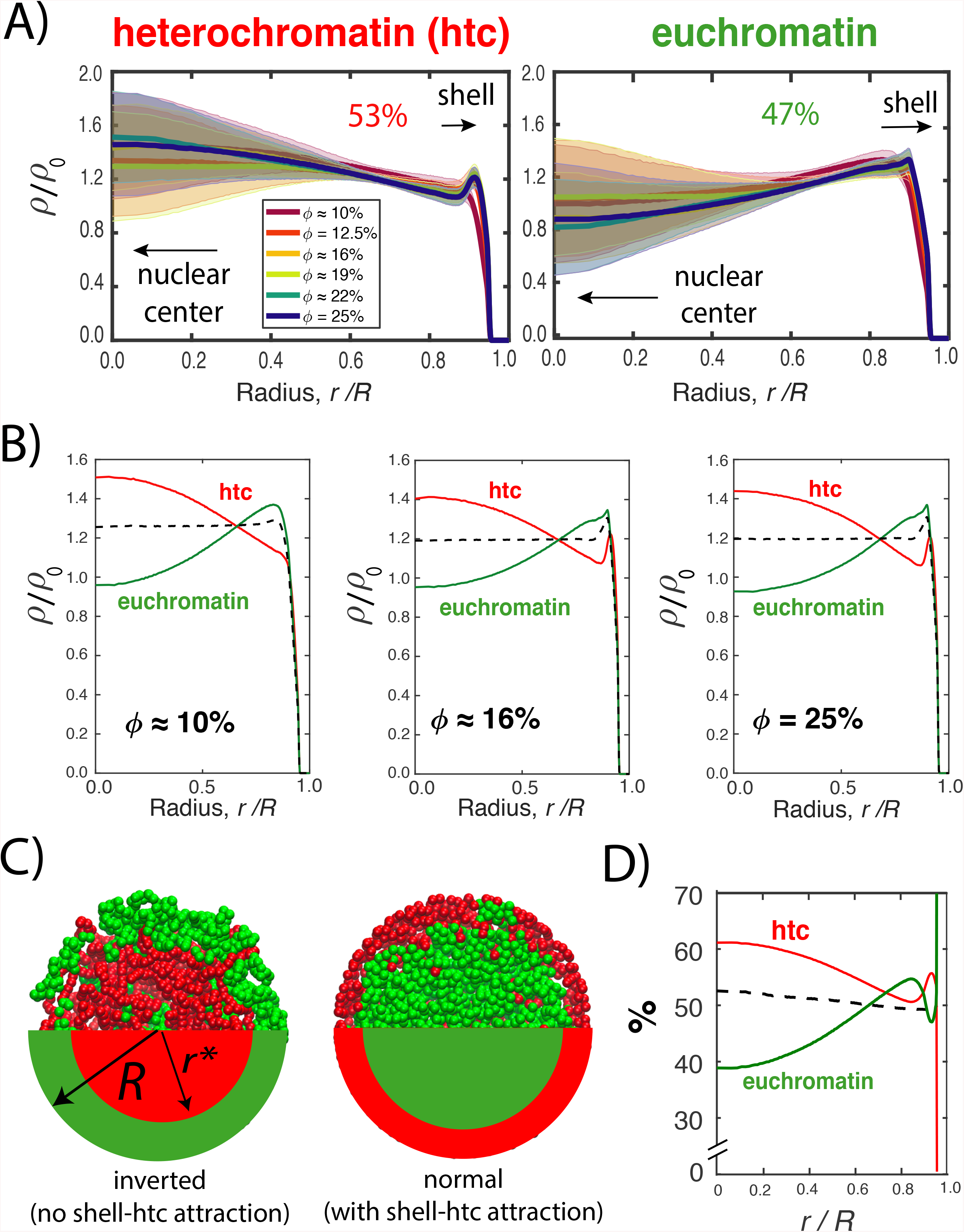
Radial concentration profiles A) The normalized radial density profiles of heterochromatin and euchromatin as a function of the rescaled radial distance from the sphere center for various volume fractions *ϕ* for a heterochromatin stiffness of *K*/*ε* = 2. The heterochromatin fraction is *f* = 0.5. The numbers indicate percentages of respective chromatin types in the central region (see text for details). B) The normalized radial density profiles shown in A for three volume fractions. Error bars as in B are not shown for clarity. Dashed curves are the control simulations with *K*/*ε* = 0 (i.e., all euchromatin). C) Arbitrary simulation snapshot showing the cross-section of the chromatin-filled shells, in which *n_ch_* = 6 chains are spherically confined. Heterochromatin and euchromatin are represented by red and green beads/shades, respectively. D) Relative occupancy of two chromatin types across the nucleus.

This inverted heterochromatin distribution, with heterochromatin at the nuclear interior, is consistent with the enhanced euchromatin concentration near the nuclear periphery (Figs. 2A and B). While heterochromatin concentration is lower than euchromatin at the nuclear boundary, heterochromatin localization exhibit a concentration-dependent behavior (Fig. 2B). A previous study has reported such concentration-dependent heterochromatin enhancement near the nuclear periphery [39]. Accordingly, above the volume fractions of *ϕ* ≈ 10%, less flexible heterochromatin homopolymer chains localize near the large curvature regions at the periphery to minimize the energy penalty against bending [39]. In Figs. 2A and B, we also observe this trend with our heteropolymer chromatin chains as the volume fraction is increased from *ϕ* ≈ 10% to *ϕ* ≈ 25% at *R*/*r* → 1. However, the peripheral heterochromatin localization at the shell does not seem to affect the concentration difference between heterochromatin and euchromatin in the nuclear interior (Fig. 2A,B).

The central heterochromatin coalescence that we quantify via concentration profiles is also evident from the representative snapshots taken from the final frames of corresponding MD simulations (Fig. 2C). The snapshot on the left-hand side of Fig. 2C demonstrates the chromatin organization in the absence of shell-heterochromatin attraction. A visual inspection reveals the systematic localization of the heterochromatin (green) towards the nuclear center and increased euchromatin (read) occupation at the nuclear periphery. This inverted chromatin organization can be compared to the snapshot from a separate simulation, for which the attraction between the shell and heterochromatin is introduced (the snapshot on the right-hand side of Fig. 2C). As we will discuss further in the next sections, the shell-chromatin attraction increases the concentration of the heterochromatin at the periphery, while enhancing euchromatin presence in the nuclear interior, as expected for a conventional nuclear organization [5, 52].

To obtain further insights into the higher concentration of heterochromatin near the nuclear center as compared to euchromatin, we calculate the relative occupancy of the two chromatin types, *n_h_*(*r*)/[*n_h_*(*r*) + *n_e_*(*r*)] and *n_e_*(*r*)/[*n_h_*(*r*)+*n_e_*(*r*)], where *n_h_*(*r*), and *n_e_*(*r*) are the number of heterochromatin and euchromatin beads, respectively, at a radial position 0 < *r* < *R*. The sum of the two relative concentrations is unity. The relative concentration profile for *ϕ* 10% in Fig. 2D confirms the trend of central heterochromatin coalescence, and the depletion of euchromatin from the interior: On average, near the center of the nucleus, the probability of a heterochromatin bead encountering another heterochromatin bead is significantly higher than a euchromatin bead, and this trend is in stark contrast with the control (i.e., *K*/*ε* = 0, dashed curve in Fig. 2D), for which there is no flexibility difference along the chromatin.

Overall, our MD simulations are able reproduce the experimentally observed central heterochromatin coalescence [23, 24] by only considering the flexibility difference between the hetero- and euchromatin blocks; when *i*) the persistance length of the euchromatin block is less than half of heterochromatin’s, and *ii*) the two chromatin types have equal nuclear concentrations (*f* = 0.5).

### Central heterochromatin localization is insensitive to relative heterochromatin content

Up to this point, our MD simulations have considered the scenario in which there is an equal amount of euchromatin and heterochromatin in the nucleus (i.e., *f* = 0.5). Under this condition, and without chromatin-shell interactions, the heterochromatin concentration is higher than euchromatin in the nuclear interior (Fig. 2). However, for various cell types or even for various phases of the same cell type, the heterochromatin content may deviate from *f* = 0.5. For instance, for fully differentiated cells, roughly 40 % of chromatin bear heterochromatin markers [18], and this number tends to be even lower for embryonic stem cells [53]. Similarly, aging cells can exhibit a time-dependent loss of heterochromatin [14]. Therefore, we considered nuclei with heterochromatin fractions of *f* < 0.5 (Fig. 1A) to study the robustness of the central coalescence phenomenon for different total nuclear heterochromatin contents. As the value of f is decreased, so does the total heterochromatin concentration relative to euchromatin’s (Fig. 3A). However, the overall trend of the concentration profiles is identical to the *f* = 0.5 case (Fig. 2); The concentration of heterochromatin is higher near the center than the periphery, and conversely, euchromatin occupation decreases in the nuclear interior (Fig. 3A). If we rescale the heterochromatin and euchromatin concentrations by their average concentrations, the general trend is that regardless of heterochromatin content, without anchoring interactions with the nuclear shell, heterochromatin concentrates near the nuclear center as opposed to euchromatin (Fig.3B).

**FIG. 3.**
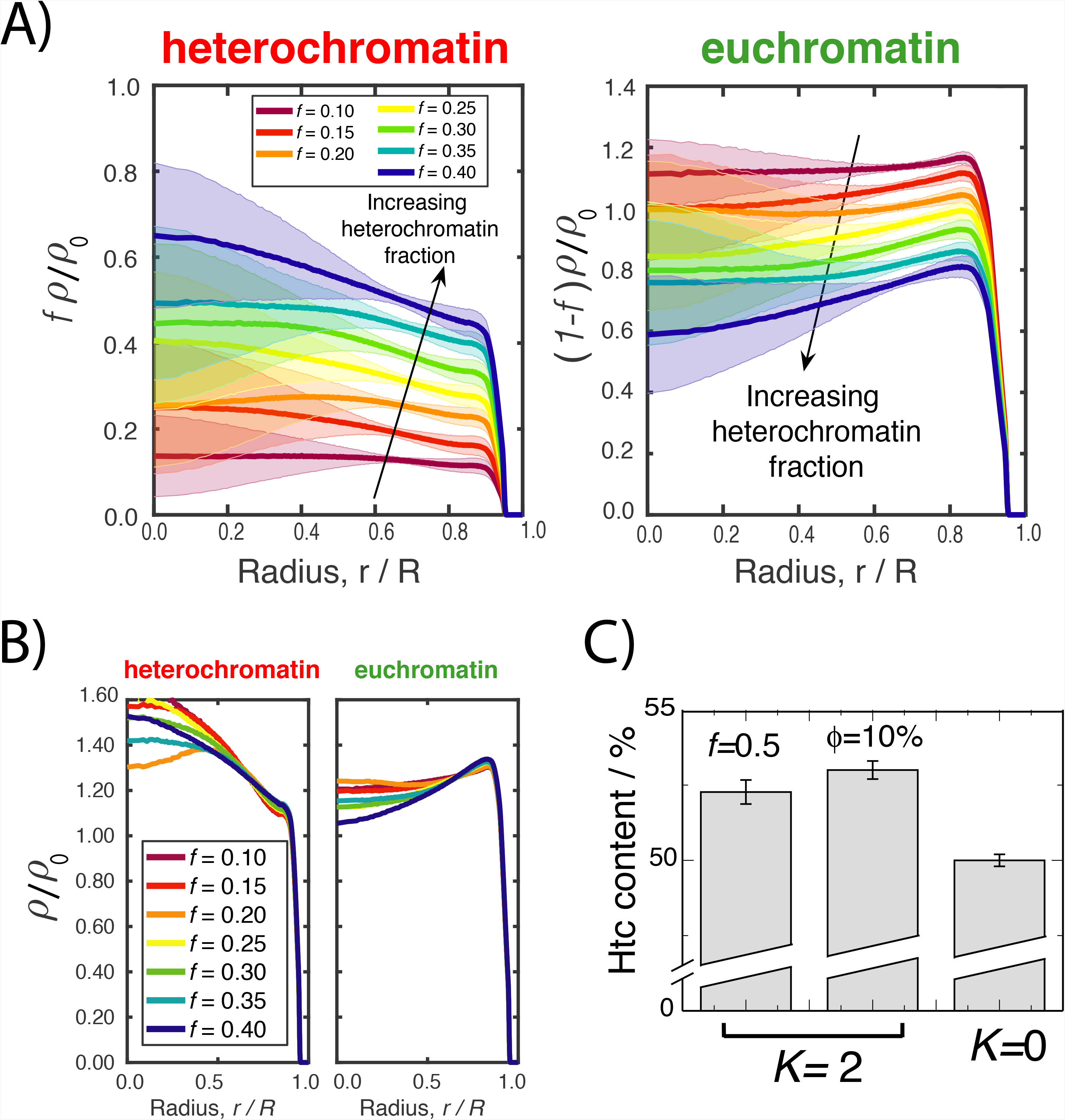
Radial concentration profiles for various heterochromatin contents *f* at a volume fraction of *ϕ* ≈ 10%. The difference in flexibility is *K* = 2*ε*/rad^2^ for all cases. The data is normalized by the rescaled concentrations to indicate the absolute difference. Arrows indicate the direction of increasing *f*. B) Concentration profiles normalized by the average concentration of respective chromatin type. Error bars are not shown for clarity. C)The percentage of heterochromatin within the central region of the nucleus averaged over various values of *ϕ* and f cases as compared to all-euchromatin case (i.e., *K*/*ε* = 0).

To obtain a quantitative measure of segregation, we calculate the cumulative heterochromatin concentration in the nuclear interior by dividing the nuclear volume into peripheral and central regions (Fig. 2C). Initially, we had defined the central region as the radius being within a range of 0 < *r* < *r*\*, where *r*\* = R/2^1/3^ ≈ 0.8 follows from the geometrical definition which splits the nucleus into two regions with equal volumes. However, this definition results in an unequal chromatin distribution in each region even for our control (i.e., *K*/*ε* = 0, all-euchromatin case) due to the depletion of monomers from the shell surface (see SI text Fig. 1). We therefore defined the value of *r*\* as the radius for which the overall chromatin concentration is equally distributed into central and peripheral regions. This criterion on average leads to a total heterochromatin accumulation in the central and peripheral regions as 53 ± 0.2% and 47 ± 0.2%, respectively, which confirms that the majority of the heterochromatin localizes in the nuclear interior (Fig.3C). While at first glance this heterochromatin accumulation in the nuclear interior seems to be weak due to the heterochromatin in the central region being only 6% higher than that in the peripheral region, this difference persists for various values of *f* and *ϕ*, and exhibits a systematic deviation from the control simulations, in which there is no flexibility difference between heterochromatin and euchromatin (Fig.3C).

### Excessive heterochromatin stiffness reduces central segregation

Our simulations with a weak bending rigidity difference (i.e., *K*/*ε* = 2) between heterochromatin and euchromatin blocks of the chromatin diblock copolymer reproduce the experimentally observed heterochromatin inversion in the absence of chromatin-NE attractions (Figs. 2 and 3). Next, to systematically investigate the effect of heterochromatin stiffness on the central coalescence, we vary the flexibility difference between *K*/*ε* = 0.5 and *K*/*ε* = 15 (see Fig. 4A), where the *K*/*ε* parameter is proportional to the persistence length of the heterochromatin (Eq. 4). Simulations in which *K*/*ε* ⪢ 1 correspond to persistance lengths on the order of the nuclear radius and are therefore less relevant to chromatin segregation, but can still be used to determine the limiting behavior. Intermediate values of *K*/*ε* can be realized experimentally by using facilitating intercalators such as propidium iodide [54, 55] or multivalency governed HP1 contacts [56].

**FIG. 4.**
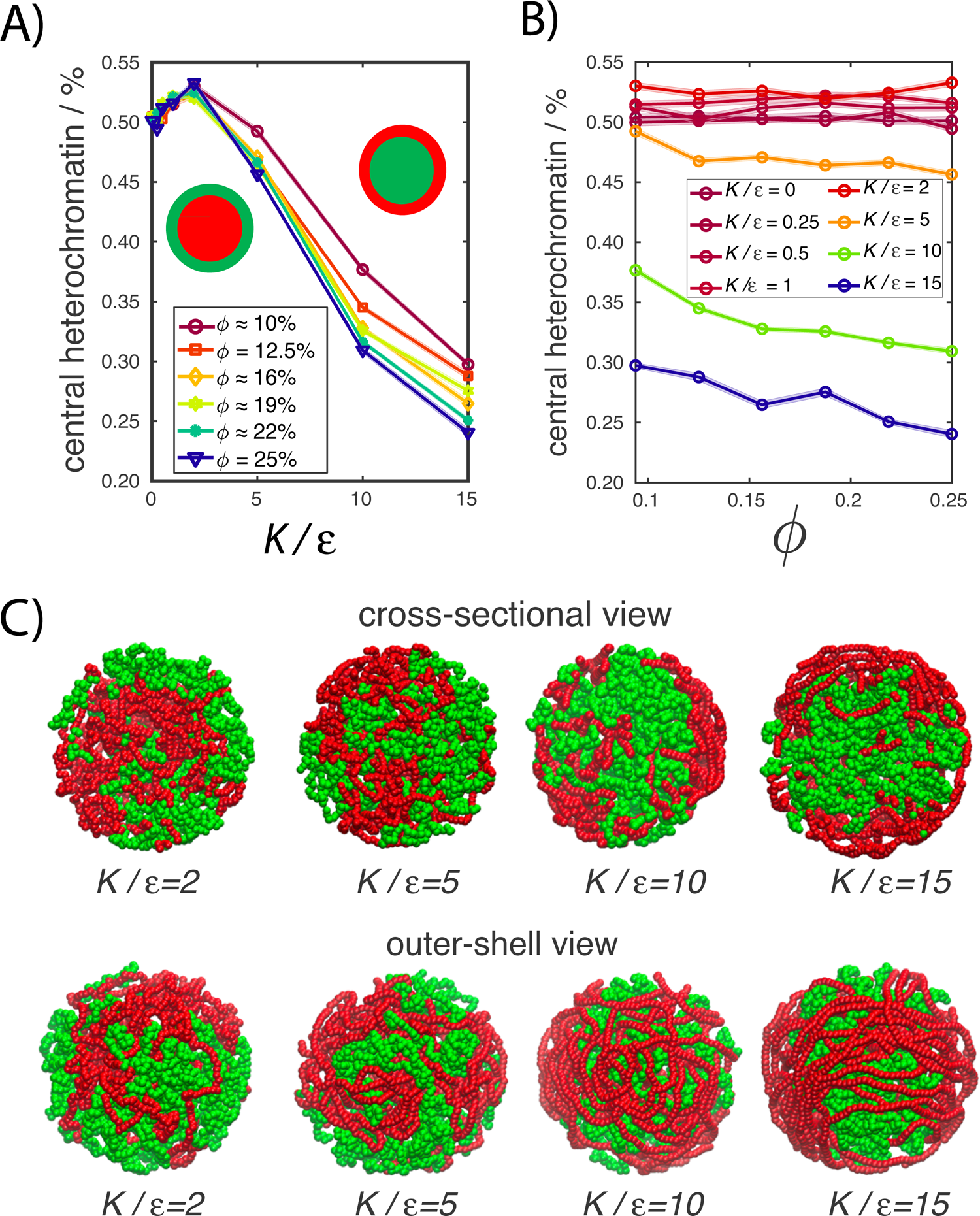
The peripheral segregation of stiffer heterochromatin A) Heterochromatin accumulated in the central region for various volume fractions as a function of the stiffness parameter. B) The same data as a function of the volume fraction but for various stiffness cases. C) Cross-sectional and outer views of spherical shells for various stiffness values. The higher the *K*/*ε* value is, the stiffer the heterochromain is. Red and green beads are heterochromatin and euchromatin beads, respectively.

Fig. 4A shows the heterochromatin accumulation in the central region as a function of *K* for various volume fractions. For weak differences in flexibility (i.e., *K*/*ε* ≈ 1), the majority of heterochromatin occupies the central region as before. As *K*/*ε* → 0, the central coalescence of heterochromatin disappears gradually. Also, for *K*/*ε* < 2, the coalescence appears to exhibit a weak dependence on the volume fraction (Fig. 5B).

**FIG. 5.**
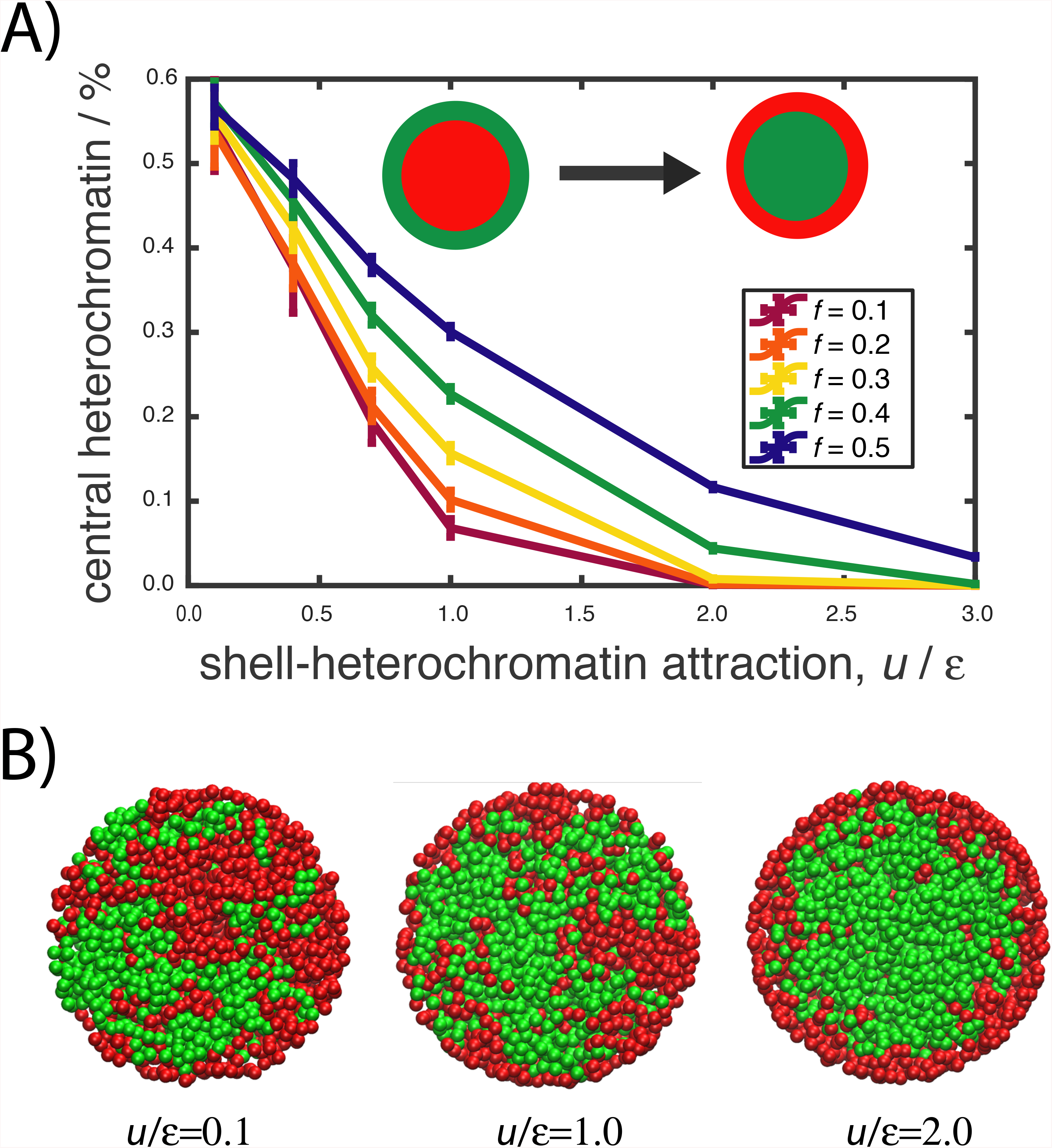
Attraction between the shell and heterochromatin decreases heterochromatin segragation in the nuclear interior A) Percentage of heterochromatin within the central region for various heterochromatin fractions ranging from *f* = 0.1 to *f* = 0.5 as a function of the shell-heterochromatin interaction strength *u*/*ε*. B) Cross-sectional views of the simulation snapshots from the respective equilibrium systems for *f* = 0.5 demonstrating heterochromatin segregation at *u*/*ε* = 0.1, 1.0, 2.0. In all cases, *ϕ* ≈ 10% and *K*/*ε* = 2.

As the strength of the bending potential, and thus the heterochromatin stiffness, is increased further (i.e., to *K*/*ε* = 5), the heterochromatin segments exhibit a more pronounced localization at the periphery (Fig. 4A,B and SI text Fig.2). At *K*/*ε* = 10, we observe the formation of circular heterochromatin loops strongly adsorbed on the inner surface of the shell (Fig. 4C). The circular segments disappear for *K*/*ε* = 15, and the nematically ordered heterochromatin segments cover the entire inner surface of the shell (Fig. 4C). Similar polymer layers were reported previously for linear semiflexible chains confined in spherical volumes in the context of confinement-induced nematic phases [57]. Overall, for biologically relevant differences in flexibility, our simulations indicate preferential heterochromatin localization in the nuclear interior.

### Attraction between shell and heterochromatin reduces central localization

After establishing the several known hallmarks of nuclear chromatin segregation with our model, next we consider heterochromatin-NE attractive interactions in more detail. In reality, these interactions are mediated by lamin-associated proteins [17, 23], which we approximate as a simple short-ranged potential in our simulations. While it oversimplifies the vast complexity of chromatin-NE interactions, it provides a measure of interaction strength needed to reverse the heterochromatin inversion. Consequences of this approximation are discussed later.

In Fig. 5A, we show the heterochromatin accumulation within the central region as a function of the attraction strength *u*/*ε* for various heterochromatin contents, *f*, for *ϕ* = 10%. As the attraction strength is increased, the heterochromatin concentration at the central region significantly decreases (Fig. 5A), and consequently the peripheral-heterochromatin concentration increases. At around an attraction strength of *u*/*ε* ≈ 0.5, the peripheral and central heterochromatin concentration are roughly 50 % (Fig. 3A). The threshold attraction strength for this behaviour is slightly weaker than the thermal energy scale 1*k_B_T*, which highlights the entropic nature of the central coalescence that we observe here. Since the attraction strength is defined per bead, the total attraction acting on a segment composed of multiple beads is stronger than 1*k_B_T*. This indicates the possibility of a cooperative behaviour, in which heterochromatin is adsorbed on the surface via a multivalent-binding mechanism [18].

As the attraction between the shell and heterochromatin is increased above the thermal energy, *u*/*ε* > 1, the fraction of surface-bound of monomers increases regardless of overall heterochromatin content (Fig. 5A,B). For the lowest amount of heterochromatin (i.e., *f* = 0.1), we observe a complete depletion of heterochromatin from the central region. For higher values of *f* (e.g., *f* = 0.5), even for relatively high attraction strengths (*u*/*ε* = 2), we observe roughly 15% of the heterochromatin in the central region (Fig. 3A,B). Notably, for *f* = 0.5, a *heterochromatin layer* is formed adjacent to periphery due to the overcrowding of the heterochromatin monomers at the inner periphery (the snapshot at the right in Fig. 5B).

Overall, our coarse-grained MD simulations show that when heterochromatin blocks of diblock chromatin polymers are less flexible, the majority of the heterochromatin localizes in the nuclear interior. This trend is non-monotonic and can be reversed either by further increasing energy penalty for heterochromatin bending (Fig. 3) or by inducing interactions between the confining shell and heterochromatin (Fig. 5).

## DISCUSSION

### Heterogeneity in chromatin flexibility can contribute to nuclear organization

Our MD simulations, where chromatin is modeled as a diblock ring copolymer, demonstrate that only a small (e.g., a factor 2 or even less) flexibility difference between the two chromatin types could be enough to segregate the majority of the heterochromatin towards the nuclear interior, away from the perimeter, if heterochromatin is weakly anchored to NE (Figs. 2,3). Importantly, this effect does not require any selective interaction between chromatin units. Our results also suggest that the central heterochromatin localization is independent of nuclear heterochromatin content (Fig. 3). Surprisingly, the central coalescence of heterochromatin is energetically very easy to overcome; Heterochromatin localizes near periphery, at its conventional position, even if NE-heterochromatin attraction is on average relatively weak (e.g., less than the thermal-energy, *k_B_T*).

The fundamental question to be discussed here with priority whether heterochromatin is less flexible than euchromatin or not. There is growing skepticism on the existence of a relatively stiffer 30-nm heterochromatin fiber *in vivo* [58]. In parallel, research into the liquid-like structure of heterochromatin-rich domains [59] reports surprising similarities with melts of flexible polymers. Hence, one may think that both chromatin types are simply mechanically indistinguishable. However, nuclear biochemical processes (e.g., histone modifications and DNA methylation) can affect the relative physical proximity between adjacent nucleosomes along the chromatin polymer by forming tightly-packed heterochromatin or loosely packed euchromatin sections. Within the heterochromatin, positional fluctuations of nucleosomes can be suppressed by neighboring nucleosomes [35, 60]. The extent of this suppression, particularly in the transverse direction of the chromatin main axis, is weaker for euchromatin as compared to heterochromatin, potentially making euchromatin effectively more flexible than heterochromatin [34], and thus, can allow the functionality of the mechanism as we demonstrate here.

The segregation mechanism that we demonstrate here could be used by cells to dynamically organize their nuclear architecture. Specifically, cells could use the suppressed flexibility of heterochromatin to save energy in processes which require re-arrangement of nuclear content. For instance, in absence of NE-chromatin attractions, heterochromatin readily segregates to the interior and promotes the formation of a single heterochromatin-rich domain [23]. This mechanism can be used as well when the cell enters mitosis, when the chromatin condenses into thicker fibers. The same mechanism can work as the cell enters mitosis, and chromatin condenses into thicker “fibers”. It is instructive to think that, with the breakage of chromatin-NE contacts followed by increasing condensation of nucleosomes, the centers of masses of the chromosomes segregate towards the nuclear interior by the mechanism described here, where sister chromatids are located prior to cell division.

There are several caveats in our simulations. One of them is that the the relaxation time of the ring polymers could exceed the simulation period that we can achieve here due to topological polymer entanglements. Biologically, high entanglement-scenario corresponds to relatively lower activity of topoisomarase II *in vivo* [61], which in fact can result in chromosomes relaxation times even longer than the duration of interphase [42]. Nevertheless, one can argue that the central heterochromatin coalescence that we demonstrate here could disappear in an ideal system where all chains are relaxed. In fact, in the simulations where non-concatenated rings are replaced by concatenated versions by allowing bond crossings, thus, eliminating entanglements [62], we observe that the amount of heterochromatin in the nuclear interior further increases (See SI text, Fig.3). Since chains can mix quicker in the absence of entanglements [41], we can conclude that the central heterochromatin coalescence shown here is not a side effect of unrelaxed chain conformations and more pronounced if topological proteins are over-expressed.

Secondly, our NE interactions assumes that heterochromatin can interact with the whole surface. In reality, there is a set of lamin-associated proteins providing such attachment [17]. Therefore, while our model over-estimates the NE-chromatin contacts, it underestimates the interaction strengths needed to keep the majority of the heterochromatin near periphery. Nevertheless, the average size of the lamin-associated domain (LADs) is as large as several million base pairs [19]. Thus, the anchoring effect of proteins on the inner surface of the NE, and resulting LADs, could be well approximated by the whole-surface-attraction scheme used here. Further, in our simulations, for a surface-attraction strengths of around ~ 1*k_B_T* per bead, an average of 10 beads are in contact with the surface (data not shown). This indicates that each heterochromatin segment is bound to the surface via ~ 10*k_B_T*, which is a reasonable value for inter-molecular interactions [63].

### Conditions for heterochromatin depletion from nuclear interior

For various nucleic acid concentrations (i.e., chromosome numbers) and for various heterochromatin contents, we systematically observe a higher concentration of heterochromatin than euchromatin in the nuclear interior (Figs. 2 and 3). Two mechanisms can reverse the central localization of heterochromatin and increase its peripheral concentration; i) An excessive bending rigidity of heterochromatin compared to euchromatin (Fig. 4), ii) attraction between heterochromatin and NE (Fig. 5). In the latter case, our simulations, which only considers non-specific attractions, clearly demonstrated how strong this effect can be even in the cases that the strength of the attraction is relatively weak. The experimentally observed negative correlation between LBR expression levels and central heterochromatin coalescence support this argument [22–24]. The former mechanism requires, in the absence of NE-chromatin attraction, a considerably high bending rigidity (i.e., the persistence length must be on the order of the nuclear dimensions) such that individual semiflexible chains enter a buckling regime [57](Fig. 4). Yet, the persistence length of heterochromatin, 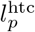, is at most several hundred nanometers [37], which is significantly lower than nuclear size. Whether other mechanisms, such as the molecular bridge formation [59] can effectively lead to sufficient stiffness still remains unknown.

### Interplay between flexibility-driven segregation and liquid-liquid phase separation

Given the complexity of the *in vivo* nuclear environment, the effect that we demonstrate here could enhance the heterochromatin-rich domain formations in the nuclear interior *in vivo* in conjunction with liquid-liquid phase separation [25, 27, 28, 64]. Chromatin with heterochromatin markers chemically favors interactions with HP1 [26, 59], and this could effectively induce an attraction between heterochromatin units. Our simulations show that these attractive interactions between heterochromatin segments may further amplify the heterochromatin inversion in the absence of NE-chromatin attraction by increasing the probability of two-body interactions between the heterochromatin. Thus, one can presume that the suppressed bending fluctuations of heterochromatin can favor the phase separation by increasing the probability of occurrence of two body interactions between heterochromatin segments, and in turn, increase the central localization of heterochromatin under weak NE-chromatin binding [23].

In conclusion, a coarse-grained polymer model considering heteropolymer nature of chromosome chains demonstrated that a weak mechanical flexibility difference along the chromatin fiber is capable of reproducing the experimental observed heterochromatin segregation regimes. Furthermore, this segregation is independent of chromatin content and relative heterochromatin concentrations and can be reversed by weak (~ *k_B_T*) shell-chromatin interactions. Yet various pieces of the puzzle are missing, and while the models like ours can help us understand large scale and long-time behavior of genome, they are unable to reveal the molecular-levels details of chromatin organization. In particular, the explicit roles of nuclear proteins, of their concentrations in cellular confinement [65], and of the deformability of the NE [66] on genomic architecture remains elusive. Elucidating the interplay of these effects on the large scale behavior of the genome will be tackled in future studies.

## ACKNOWLEDGMENTS

This work was supported by the Alexander von Humboldt-stiftung (AvH), and Scientific and Technological Research Council of Turkey (TUBITAK) via 2232 project (2018/4), 119C010. We acknowledge use of computational resources from the Max-Planck Computational and Data Facilities (MPCDF).

## References

[1] R. D. Kornberg, Science 184, 868 (1974).

[2] S. I. S. Grewal and S. C. R. Elgin, Nature 447, 399 (2007).

[3] N. Gilbert, S. Boyle, H. Fiegler, K. Woodfine, N. P. Carter, and W. A. Bickmore, Cell 118, 555 (2004).

[4] B. Harmon and J. Sedat, PLOS Biology 3, e67 (2005).

[5] M. R. Branco and A. Pombo, Trends in Cell Biology 17, 127 (2007).

[6] T. Jenuwein and C. D. Allis, Science 293, 1074 (2001).

[7] Cell 135, 9 (2008).

[8] D. Zink, A. H. Fische, and J. A. Nickerson, Nature Reviews Cancer 4, 677 (2004).

[9] D. M. Carone and J. B. Lawrence, Seminars in Cancer Biology 23, 99 (2013).

[10] P. Taimen, K. Pfleghaar, T. Shimi, D. Möller, K. Ben-Harush, M. R. Erdos, S. A. Adam, H. Herrmann, O. Medalia, F. S. Collins, A. E. Goldman, and R. D. Goldman, Proceedings of the National Academy of Sciences 106, 20788 (2009).

[11] K. Bercht Pfleghaar, P. Taimen, V. Butin-Israeli, T. Shimi, S. Langer-Freitag, Y. Markaki, A. E. Goldman, M. Wehnert, and R. D. Goldman, Nucleus 6, 66 (2015).

[12] D. K. Shumaker, T. Dechat, A. Kohlmaier, S. A. Adam, M. R. Bozovsky, M. R. Erdos, M. Eriksson, A. E. Goldman, S. Khuon, F. S. Collins, T. Jenuwein, and R. D. Goldman, Proc Natl Acad Sci USA 103, 8703 (2006).

[13] E. Haithcock, Y. Dayani, E. Neufeld, A. J. Zahand, N. Feinstein, A. Mattout, Y. Gruenbaum, and J. Liu, Proceedings of the National Academy of Sciences of the United States of America 102, 16690 (2005).

[14] R. Yu, B. McCauley, and W. Dang, Human Genetics, 1 (2020).

[15] D. M. Gilbert, The Journal of Cell Biology 152, F11 (2001).

[16] A. Akhtar and S. M. Gasser, Nature Reviews Genetics 8, 507 (2007).

[17] Y. Y. Shevelyov and S. V. Ulianov, Cells 8, 136 (2019).

[18] B. van Steensel and A. S. Belmont, Cell 169, 780 (2017).

[19] L. Guelen, L. Pagie, E. Brasset, W. Meuleman, M. B. Faza, W. Talhout, B. H. Eussen, A. de Klein, L. Wessels, W. de Laat, and B. van Steensel, Nature 453, 948 (2008).

[20] J. C. Harr, T. R. Luperchio, X. Wong, E. Cohen, S. J. Wheelan, and K. L. Reddy, The Journal of Cell Biology 208, 33 (2015).

[21] M. Zwerger, C. Y. Ho, and J. Lammerding, Annual Review of Biomedical Engineering 13, 397 (2011).

[22] A. L. Olins, G. Rhodes, D. B. M. Welch, M. Zwerger, and D. E. Olins, Nucleus 1, 53 (2010).

[23] I. Solovei, A. S. Wang, K. Thanisch, C. S. Schmidt, S. Krebs, M. Zwerger, T. V. Cohen, D. Devys, R. Foisner, L. Peichl, H. Herrmann, H. Blum, D. Engelkamp, C. L. Stewart, H. Leonhardt, and B. Joffe, Cell 152, 584 (2013).

[24] I. Solovei, M. Kreysing, C. LanctOt, S. KOsem, L. Peichl, T. Cremer, J. Guck, and B. Joffe, Cell 137, 356 (2009).

[25] B. A. Gibson, L. K. Doolittle, L. E. Jensen, N. Gamarra, S. Redding, and M. K. Rosen, bioRxiv, 523662 (2019).

[26] Q. MacPherson, B. Beltran, and A. J. Spakowitz, Proceedings of the National Academy of Sciences 115, 12739 (2018).

[27] A. R. Strom, A. V. Emelyanov, M. Mir, D. V. Fyodorov, X. Darzacq, and G. H. Karpen, Nature 547, 241 (2017).

[28] A. R. Strom, R. J. Biggs, E. J. Banigan, X. Wang, K. Chiu, C. Herman, J. Collado, F. Yue, J. C. R. Politz, L. J. Tait, D. Scalzo, A. Telling, M. Groudine, C. P. Brangwynne, J. F. Marko, and A. D. Stephens, bioRxiv (2020), 10.1101/2020.10.09.331900.

[29] L. Leibler, Macromolecules (1980).

[30] C. E. Sing, J. W. Zwanikken, and M. Olvera de la Cruz, Nature Materials 13, 694 (2014).

[31] C. Singh, M. Goulian, A. J. Liu, and G. H. Fredrickson, Macromolecules 27, 2974 (1994).

[32] F. S. Bates and G. Fredrickson, Macromolecules 27, 1065 (1993).

[33] S. Mao, Q. MacPherson, J. Qin, and A. J. Spakowitz, Soft Matter 13, 2760 (2017).

[34] N. Kepper, D. Foethke, R. Stehr, G. Wedemann, and K. Rippe, Biophysical Journal 95, 3692 (2008).

[35] O. Müller, N. Kepper, R. Schöpflin, R. Ettig, K. Rippe, and G. Wedemann, Biophysical Journal 107, 2141 (2014).

[36] M. A. Ricci, C. Manzo, M. F. García-Parajo, M. Lakadamyali, and M. P. Cosma, Cell 160, 1145 (2015).

[37] K. Rippe, Trends in Biochemical Sciences 26, 733 (2001).

[38] J. Langowski, The European Physical Journal E 19, 241 (2006).

[39] P. R. Cook and D. Marenduzzo, The Journal of Cell Biology 186, 825 (2009).

[40] L. A. Mirny, Chromosome Research 19, 37 (2011).

[41] J. D. Halverson, J. Smrek, K. Kremer, and A. Y. Grosberg, Reports on Progress in Physics 77, 022601 (2014).

[42] A. Rosa and R. Everaers, PLoS Computational Biology 4, e1000153 (2008).

[43] K. Kremer and G. S. Grest, Journal of Chemical Physics 92, 5057 (1998).

[44] S. de Nooijer, J. Wellink, B. Mulder, and T. Bisseling, Nucleic Acids Research 37, 3558 (2009).

[45] M. J. Stevens, Biophysical Journal 80, 130 (2001).

[46] J. A. Anderson, C. D. Lorenz, and A. Travesset, Journal of Computational Physics 227, 5342 (2008).

[47] J. Glaser, T. D. Nguyen, J. A. Anderson, P. Lui, F. Spiga, J. A. Millan, D. C. Morse, and S. C. Glotzer, Computer Physics Communications 192, 97 (2015).

[48] M. Girard, A. Ehlen, A. Shakya, T. Bereau, and M. O. de la Cruz, Computational Materials Science 167, 25 (2019).

[49] A. Stukowski, Modelling and Simulation in Materials Science and Engineering 18, 015012 (2009).

[50] W. Humphrey, A. Dalke, and K. Schulten, Journal of molecular graphics 14, 33 (1996).

[51] J. D. Halverson, W. B. Lee, G. S. Grest, A. Y. Grosberg, and K. Kremer, The Journal of Chemical Physics 134, 204905 (2011).

[52] H. Dehghani, G. Dellaire, and D. P. Bazett-Jones, Micron (Oxford, England: 1993) 36, 95 (2005).

[53] S. Efroni, R. Duttagupta, J. Cheng, H. Dehghani, D. J. Hoeppner, C. Dash, D. P. Bazett-Jones, S. Le Grice, R. D. G. McKay, K. H. Buetow, T. R. Gingeras, T. Misteli, and E. Meshorer, Cell stem cell 2, 437 (2008).

[54] I. D. Vladescu, M. J. McCauley, M. E. Nuñez, I. Rouzina, and M. C. Williams, Nature Methods 4, 517 (2007).

[55] A. D. Stephens, E. J. Banigan, S. A. Adam, R. D. Goldman, and J. F. Marko, Molecular biology of the cell 28, 1984 (2017).

[56] S. Kilic, A. L. Bachmann, L. C. Bryan, and B. Fierz, Nature Communications 6, 1 (2015).

[57] A. Nikoubashman, D. A. Vega, K. Binder, and A. Milchev, Physical Review Letters 118, 217803 (2017).

[58] E. Fussner, R. W. Ching, and D. P. Bazett-Jones, Trends in Biochemical Sciences 36, 1 (2011).

[59] K. Hiragami-Hamada, S. Soeroes, M. Nikolov, B. Wilkins, S. Kreuz, C. Chen, I. A. De La Rosa-Velázquez, H. M. Zenn, N. Kost, W. Pohl, A. Chernev, D. Schwarzer, T. Jenuwein, M. Lorincz, B. Zimmermann, P. J. Walla, H. Neumann, T. Baubec, H. Urlaub, and W. Fischle, Nature Communications 7, 11310 (2016).

[60] S. S. Ashwin, T. Nozaki, K. M. P. o. the, and 2019, Proceedings of the National Academy of Sciences 116, 19939 (2019).

[61] S. M. Vos, E. M. Tretter, B. H. Schmidt, and J. M. Berger, Nature Publishing Group 12, 1 (2011).

[62] J. Farago, H. Meyer, J. Baschnagel, and A. N. Semenov, Physical review. E, Statistical, nonlinear, and soft matter physics 85(2012).

[63] J. N. Israelachvili, Intermolecular and Surface Forces, Revised Third Edition (Academic Press, 2011).

[64] C. P. Brangwynne, P. Tompa, and R. V. Pappu, Nature Physics 11, 899 (2015).

[65] A. Erbaş, M. O. de la Cruz, and J. F. Marko, Biophysical Journal 116, 1609 (2019).

[66] S. Seirin-Lee, F. Osakada, J. Takeda, S. Tashiro, R. Kobayashi, T. Yamamoto, and H. Ochiai, PLoS Computational Biology 15, e1007289 (2019).

