## Supplementary Information for "Heterochromatin flexibility contributes to chromosome segregation in the cell nucleus"

Martin Girard

*Max-Planck Institute for Polymer Research, 55128 Mainz, Germany*

Monica Olvera de la Cruz

*Department of Materials Science and Engineering, Department of Chemistry, Department of Chemical and Biological Engineering and Department of Physics, Northwestern University, Evanston, Illinois 60208, USA*

John F. Marko

*Department of Physics & Astronomy and Department of Molecular Biosciences, Northwestern University, Evanston, Illinois 60208, USA*

Aykut Erbaş

*UNAM-National Nanotechnology Research Center and Institute of Materials Science & Nanotechnology, Bilkent University, Ankara 06800, Turkey*

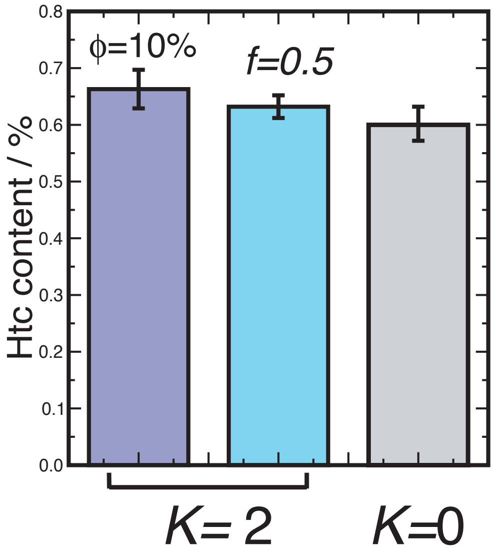


FIG. S1. The accumulation percentage of heterochromatin in the central region of nuclear volume averaged over various volume fractions $\phi$ and heterochromatin content, $f$. To separate the central and peripheral regions, the geometrical definition $r^* = R/2^{1/3} \approx 0.8$ is used. Notice that even for the control simulations with $K/\varepsilon = 0$, more than 50 % of the heterochromatin is located in the central region. The units of $K$ is $\varepsilon/rad^2$.

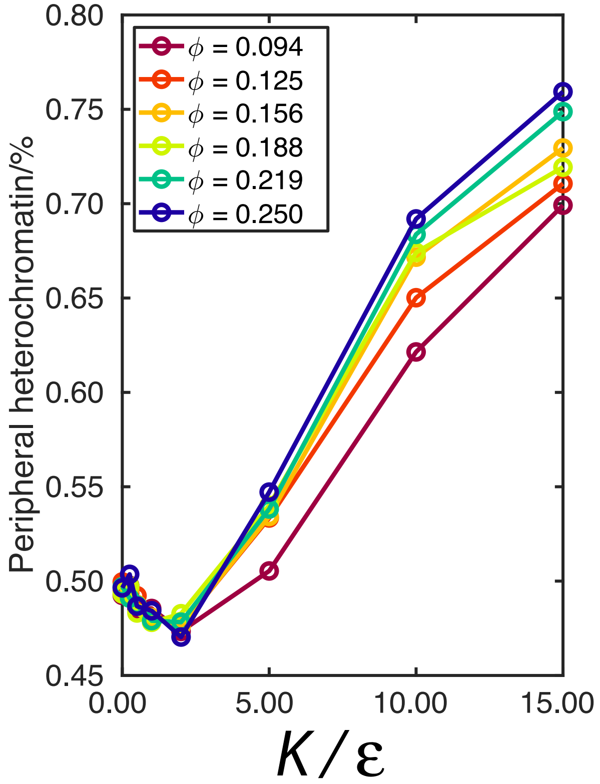


FIG. S2. The accumulation percentage of heterochromatin in the peripheral region of the nuclear volume for various volume fractions. To separate the central and peripheral regions, the definition given in the main text is used.

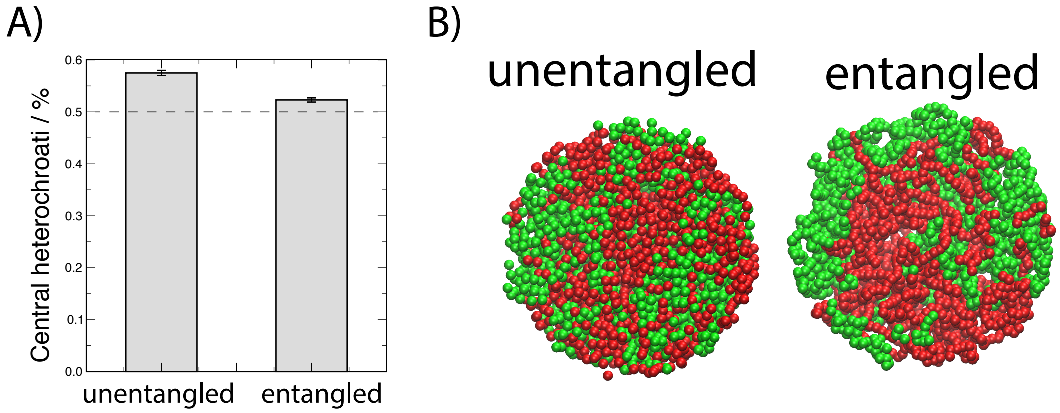


FIG. S3. Effect of entanglements on heterochromatin coalescence for $f = 0.5$ and $\phi = 10\%$. The unentangled systems biologically corresponds to high topo II protein activity in interphase. The snapshot on the right shows the distribution of heterochromatin (red) and euchromatin (green) monomers in a system that bond can cross and the one on the right shows the case that bond cannot cross. In the middle, percentage of heterochromatin monomers for the two cases in the central region is shown. To obtain unentangled polymer systems, FENE bond in the main simulations (see Eq. 2 in the main text) is replaced by a harmonic bond potential $U_H = K(r - 1.5\sigma)^2$ to allow bond crossing, where the bond strength is $80\varepsilon/\sigma^2$. To separate the central and peripheral regions, the definition given in the main text is used.
